# CryoCodex: learning discrete structural representations for cryo-EM map post-processing

**DOI:** 10.64898/2026.09.30.755675

**Authors:** Bin Cheng, Baoquan Su, Jianyi Yang

**Affiliations:** MOE Frontiers Science Center for Nonlinear Expectations, Research Center for Mathematics and Interdisciplinary Sciences, Shandong University, Qingdao, China

## Abstract

Cryo-EM maps frequently suffer from weak structural signals, spatially variable quality, and noise, complicating interpretation and downstream modeling. Existing post-processing approaches treat density enhancement, quality assessment, and masking as separate tasks, often amplifying residual noise or introducing local artifacts. Here, we present CryoCodex, a unified framework that jointly performs density enhancement, local quality estimation, and molecular masking. CryoCodex introduces a multi-head, multi-scale vector-quantization module that projects encoder-derived latent features onto a shared codebook of structural prototypes. By imposing discrete constraints across feature subspaces and spatial scales, the codebook suppresses stochastic noise while preserving recurrent structural patterns. On a non-redundant test set of 126 experimental maps spanning 3–8 Å, CryoCodex outperformed four representative methods, achieving a mean map–model FSC-0.5 of 3.93 Å versus 4.62 Å for EMReady2 and 6.87 Å for the deposited maps. Downstream automated *de novo* model building with Phenix further confirmed the quality of the CryoCodex-enhanced maps. These gains extended to maps containing nucleic acids, where CryoCodex produced the best map–model agreement and improved model building. Half-map cross-validation indicated that these improvements are anchored to experimental data rather than driven by half-map-specific noise or structural hallucination. CryoCodex also delivered reliable local quality estimates and molecular masks within the same framework. Together, these results establish CryoCodex as a practical tool for cryo-EM map post-processing and structural determination in challenging cases.

## Introduction

Cryo-electron microscopy (cryo-EM) has become an indispensable tool in determining the three-dimensional structures of biological macromolecules at near-atomic resolution. With continuous advances in data acquisition technologies and image processing algorithms, the number of cryo-EM entries deposited in the Electron Microscopy Data Bank (EMDB)^1^ has increased rapidly. However, owing to factors such as radiation damage, molecular motion, and reconstruction errors, experimental density maps can still exhibit limited contrast, background density fluctuations, and pronounced heterogeneity in local resolution across the molecular structure^2–5^. These limitations can obscure structural features and substantially increase the difficulty of subsequent structure determination and atomic model building^6–10^.

To improve the visibility and interpretability of these structural features, various post-processing methods have been developed to enhance reconstructed cryo-EM density maps. In recent years, deep learning-based approaches^11–14^ have shown substantial advantages over conventional sharpening methods^15–20^ by directly learning nonlinear transformations in real space. Representative methods such as EMReady^11^ and EMReady2^13^ have demonstrated strong capabilities in suppressing background noise and enhancing structural density. More recently, physics-informed deep learning approaches, such as LocScale-2.0^21^, have also emerged. However, robust enhancement remains challenging in regions where structurally meaningful signals are weak and approach the level of surrounding density fluctuations. Highly flexible continuous feature representations are advantageous for recovering complex structural information, but such flexibility may also increase the tendency to fit sample-specific fluctuations or unstable local patterns when the experimental signal is limited. Conversely, overly aggressive suppression of such variations may remove weak yet genuine structural features. A central design challenge is therefore to constrain enhancement while retaining the flexibility needed to represent diverse structural features.

A further challenge is that the effects of map enhancement can vary substantially across a cryo-EM map. Different regions within the same cryo-EM map may differ markedly in density quality and modelability^2, 22–24^, and improvement in a global metric does not necessarily imply comparable enhancement across all local regions. Most existing deep learning-based enhancement methods primarily provide an enhanced density map, offering limited information about the quality of individual regions after processing. This is particularly important for regions of weak density, where visually sharpened features may not necessarily correspond to reliable structural information. A spatially resolved quality estimate associated with the enhanced map can therefore indicate not only where density has been improved, but also where the resulting features can be interpreted with greater confidence.

Here we present CryoCodex, a deep-learning framework that combines map enhancement with local quality estimation and molecular masking. Its multi-head, multi-scale vector-quantization (MMVQ) module represents latent features using a learned codebook across feature subspaces and spatial scales. Coarse-to-fine residual quantization imposes discrete constraints on the representation, while gated fusion retains continuous information. Together, these components are designed to reduce the tendency to fit unstable density fluctuations while retaining the flexibility to represent structural detail. The network jointly predicts enhanced density, voxel-wise quality estimates for the input and enhanced maps, and a molecular mask. Assessment on a held-out benchmark of 126 experimental maps demonstrates the superior performance of CryoCodex for all considered tasks.

## Results

### Overview of CryoCodex

Figure 1 illustrates the overall framework of CryoCodex. To ensure computational feasibility, the input cryo-EM map is first partitioned into overlapping density boxes for network inference. Each density box is processed by MMVQ-UNet, which simultaneously predicts four-channel outputs: the enhanced density box, the local quality of the input density, the local quality of the enhanced density, and a molecular-region mask. The overlapping box-wise predictions are then assembled to generate the corresponding four-channel full-volume outputs. Conceptually, MMVQ-UNet can be viewed as a U-Net architecture in which the continuous latent representations generated by the network are denoised through a multi-head, multi-scale vector-quantization (MMVQ) module.

The U-Net architecture was adapted from the recurrent U-Net (RU-Net) developed in CryoAtom2 ^7^. Inspired by the ability of the RU-Net in CryoAtom2 to identify key atoms, we reasoned that its combination of convolutional operations and global attention enables effective extraction of fine-grained, atomic-level features, making it well suited for cryo-EM density enhancement. The network consists primarily of an encoder and a decoder. The encoder progressively downsamples the input density map to extract higher-level semantic information, represented as continuous features in the latent space, whereas the decoder upsamples these latent features to restore the original spatial resolution. Between the encoder and decoder, we introduce a specifically designed multi-head, multi-scale vector-quantization (MMVQ) module. The use of vector quantization (VQ) is motivated by the view that latent features in cryo-EM maps comprise recurrent structural patterns corrupted by stochastic noise^25^. By projecting continuous features onto a discrete codebook of representative prototypes, VQ suppresses noise while preserving structurally informative features. The MMVQ module is described in detail below.

The MMVQ module is designed to perform denoising in the density enhancement task (Fig. 1a). The term “multi-head” refers to partitioning the continuous latent features produced by the encoder into multiple feature subspaces, a design inspired by the success of multi-head attention in Transformers. “Multi-scale” refers to quantizing the continuous features within each subspace across multiple spatial resolutions, using representative prototypes drawn from a shared discrete codebook. The multi-scale quantization process is illustrated in Fig. 1b, with further details provided in the Methods. By introducing discrete constraints across both feature subspaces and spatial scales, MMVQ provides a structured constraint on latent feature learning. The quantized representation is subsequently combined with the continuous features through a learnable gated fusion before decoding. The resulting shared representation is then used for density enhancement, local quality assessment, and molecular region prediction within a unified framework.

**Fig. 1.**
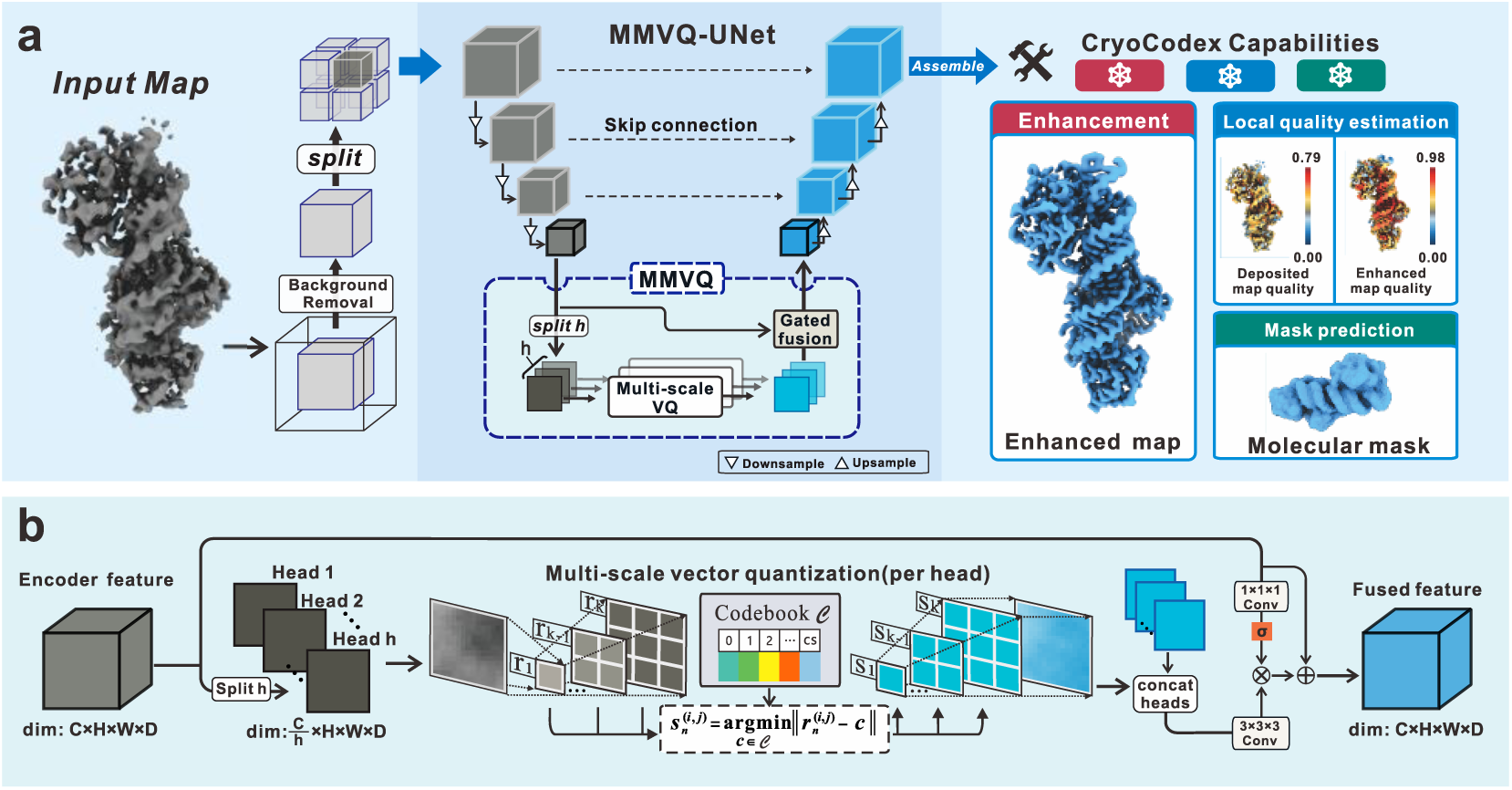
| CryoCodex framework. **a** The input density map is divided into fixed-size density patches (48 × 48 × 48 voxels) and processed by MMVQ-UNet, which combines multi-head, multi-scale vector-quantization (MMVQ) with gated fusion of quantized and continuous features. The network produces four-channel output patches, which are then assembled into an enhanced density map, local quality maps for the input and enhanced maps (in-score and out-score, respectively), and a molecular mask. **b** Schematic illustration of multi-scale vector quantization within a single quantization head. Features are quantized at multiple spatial scales through nearest-neighbor matching with a shared codebook to obtain discrete representations at different scales.

### CryoCodex outperforms existing methods in map quality enhancement

We first evaluated CryoCodex on a test set of 126 experimental cryo-EM maps from the EMDB^1^. These maps span reported resolutions from 3 to 8 Å and cover a broad range of structural classes, including proteins, nucleic acids, and protein–nucleic acid complexes. Importantly, the test set was filtered against the training set to remove redundancy and prevent evaluation bias from memorization (see Methods).

To evaluate the performance of CryoCodex, we compared it with the deposited maps and four representative deep-learning-based cryo-EM map post-processing methods: CryoTEN^12^, DEMO-EMReF^14^, EMReady^11^, and EMReady2^13^. Map quality was assessed using two complementary metrics: the unmasked map–model FSC at the 0.5 cutoff (hereafter referred to as FSC-0.5 unless otherwise specified), calculated using phenix.mtriage^26^, and the Q-score^27^, implemented in the MapQ plugin of UCSF Chimera^28^. FSC-0.5 reflects global agreement between the density map and the corresponding atomic model, whereas Q-score measures local map resolvability around atomic positions. Detailed definitions of all metrics are provided in the Methods.

FSC-0.5 comparisons were restricted to a common subset of 120 targets for which all evaluated methods produced valid results (i.e., successful phenix.mtriage runs). As summarized in Supplementary Table 1, CryoCodex achieved the lowest mean FSC-0.5 (3.93 Å), outperforming EMReady2 (4.62 Å) and DEMO-EMReF (4.82 Å). In addition, CryoCodex improved the FSC-0.5 value by more than 1 Å relative to the deposited maps in 75.83% of cases, compared with 43.33% for EMReady2 and 45.00% for DEMO-EMReF. Supplementary Data 1 provides the complete per-target FSC-0.5, and Supplementary Fig. 1 shows that FSC-0.5 improvements did not exhibit a clear monotonic decline with decreasing resolution.

For Q-score, valid results were available for all evaluated methods on all 126 maps; the complete test set was therefore used for this comparison. As shown in Supplementary Table 1 and Fig. 2b,c, CryoCodex achieved the highest mean Q-score of 0.503, compared with 0.460 for EMReady2 and 0.422 for the deposited maps. CryoCodex improved the Q-score by >0.05 in 73.8% of cases (vs. 35.7% for EMReady2) and by >0.08 in 57.9% of cases (vs. 11.1% for EMReady2). Notably, the scatter plots show that CryoCodex yields larger improvements for maps with higher deposited Q-scores, relative to both EMReady2-enhanced and deposited maps. This trend is consistent with the role of the MMVQ module: when the underlying structural signal is stronger, as reflected by a higher deposited Q-score, MMVQ more effectively maps the latent representation onto structurally relevant codebook prototypes while suppressing stochastic noise. Unlike FSC-0.5 improvements, Q-score gains declined progressively and became limited beyond approximately 5 Å (Supplementary Fig. 1).

**Fig. 2.**
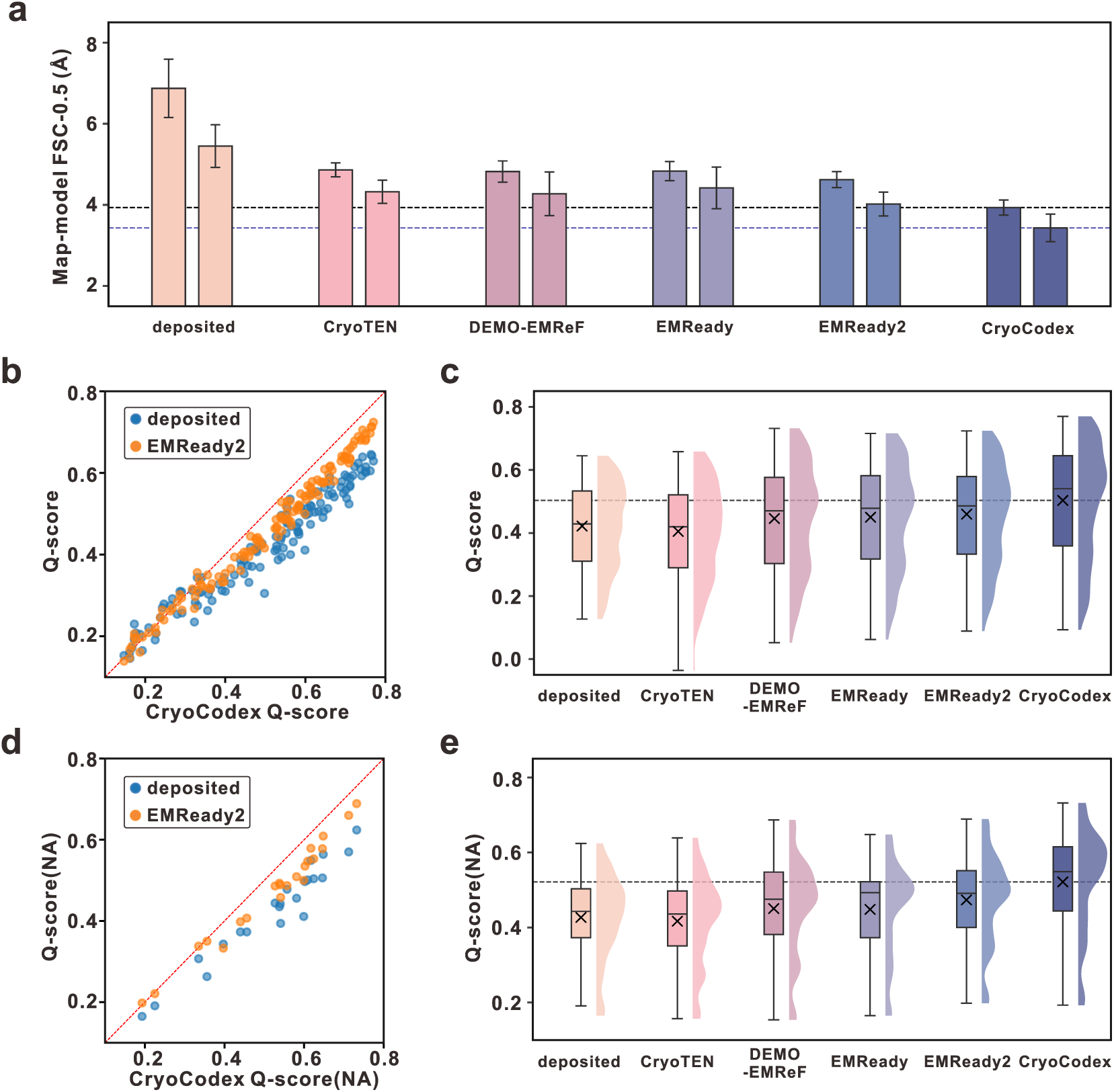
| Comparison of map–model FSC-0.5 and Q-score across map enhancement methods. **a** Map-model FSC-0.5 on the common subsets with valid results for all evaluated methods for the overall benchmark (n = 120; first bar) and the nucleic-acid subset (n = 20; second bar). The black and blue dashed lines indicate the mean CryoCodex performance on the overall and nucleic-acid subsets, respectively. Error bars represent the standard error of the mean (SEM). **b, d** Q-score comparison between CryoCodex and deposited maps or EMReady2 on the full test set (**b**) and nucleic-acid subset (**d**). **c, e** Q-score distributions across all methods on the full test set (**c**) and nucleic-acid subset (**e**). Lower FSC-0.5 and higher Q-scores indicate improved map quality and atomic resolvability. In (**c)** and (**e)**, boxes indicate the interquartile range, center lines indicate the median, whiskers extend to the most extreme data points within 1.5 times the interquartile range, and × symbols indicate the mean.

Fig. 3a–c illustrate a representative example of map enhancement for EMD-10387, corresponding to the inactive monomeric Glu494Ala mutant of *Achromobacter xylosoxidans* quinol-dependent nitric oxide reductase (AxqNOR; PDB ID: 6T6V). At a lower contour level (Fig. 3a), the deposited map exhibits substantial diffuse density surrounding the transmembrane region. Diffuse peripheral density, likely associated with the detergent environment, is also observed together with scattered low-level background density. CryoCodex substantially suppresses both the peripheral non-protein density and the scattered background density, resulting in a cleaner molecular envelope and clearer separation of the protein density from its surroundings. At a higher contour level (Fig. 3b), several transmembrane helical regions in the deposited map exhibit relatively weak or discontinuous density, whereas the CryoCodex-processed map shows more continuous and well-defined density along the modeled α-helices. The corresponding unmasked map-model FSC curves (Fig. 3c) further show increased agreement with the reference atomic model over a broad range of spatial frequencies, with the largest improvement observed at intermediate and higher spatial frequencies. Consistent with these observations, the FSC-0.5 resolution improved from 6.18 Å to 3.22 Å, while the Q-score increased from 0.387 to 0.532.

**Fig. 3.**
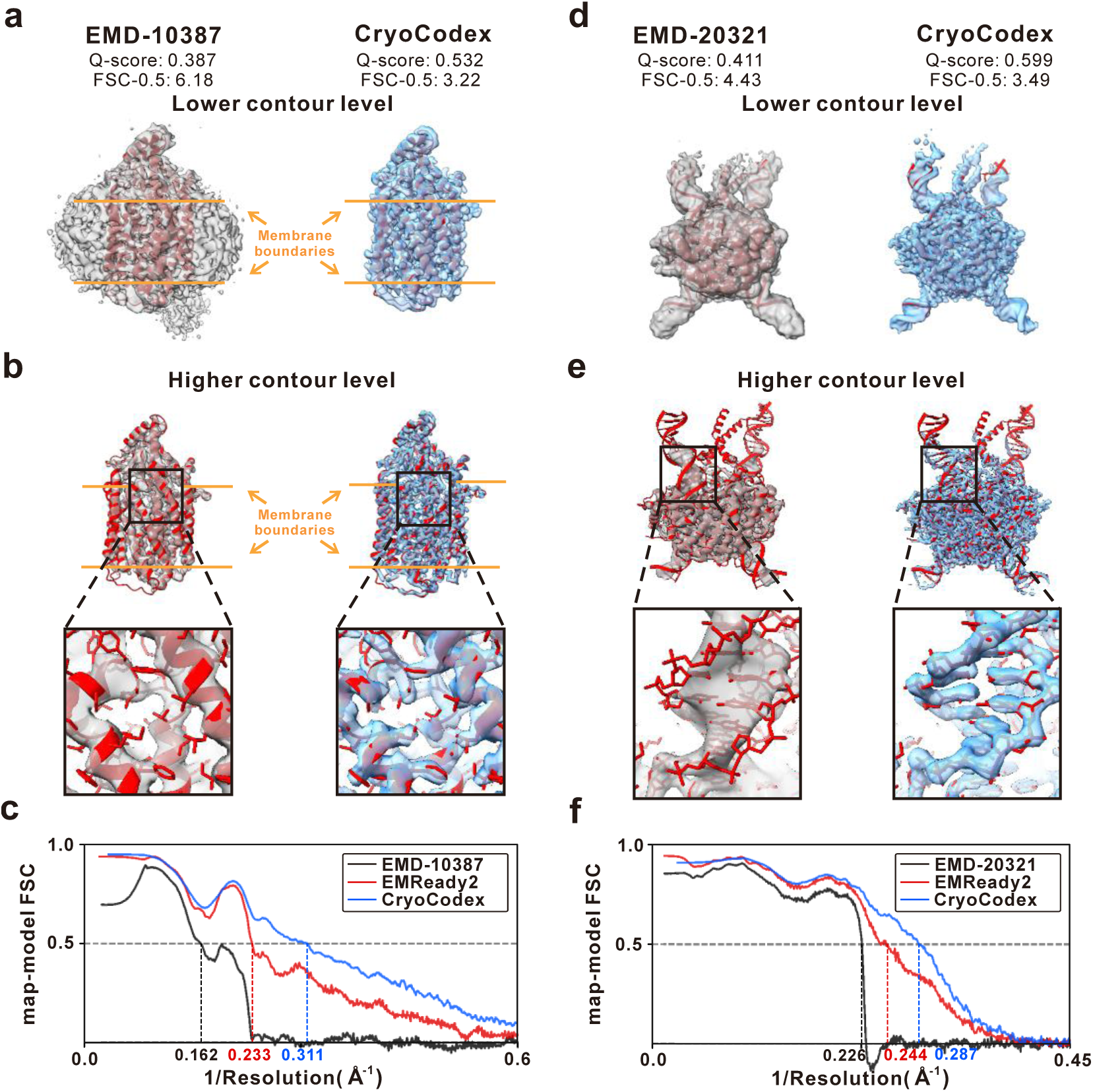
| Enhancement of protein and DNA density by CryoCodex. **a, b** EMD-10387 (PDB 6T6V; reported resolution, 4.5 Å) at lower (**a**) and higher (**b**) contour levels, with an enlarged α-helical region. The orange solid lines indicate the membrane boundaries. **c** Map-model Fourier shell correlation versus inverse resolution for EMD-10387. **d, e** EMD-20321 (PDB 6PE2; reported resolution, 4.0 Å) at lower (**d**) and higher (**e**) contour levels, with an enlarged DNA region. **f** Map-model Fourier shell correlation versus inverse resolution for EMD-20321.

### CryoCodex-enhanced maps improve downstream model building

To assess whether improvements in map quality translate into improved structural interpretability, we further evaluated downstream automated *de novo* model building. To avoid introducing additional learned priors from other deep-learning-based model-building methods, we used phenix.map_to_model^26^ as an independent automated model-building procedure. Chain-specific local density maps were extracted for all 771 chains in the test set, and atomic models were automatically built from both the deposited and processed maps. The resulting models were compared with the corresponding reference PDB^29^ structures using phenix.chain_comparison. Model-building performance was quantified using residue coverage and sequence recall (see Methods). Residue coverage measures the fraction of reference residues that are spatially matched by the built model within 3.0 Å, whereas sequence recall applies the additional requirement that spatially matched residues have the correct residue identity.

For a fair comparison across methods, statistical summaries were calculated on the common subset of 761 chains for which valid model-building results were obtained for the deposited maps and all evaluated methods; this subset included 102 nucleic acid chains. The benchmark results are summarized in Fig. 4a. CryoCodex achieved the highest mean residue coverage of 72.1%, compared with 66.7% for the second-best method, EMReady2, and 51.5% for the deposited maps. This improvement indicates that the CryoCodex-processed maps provide density that is more amenable to complete automated tracing. CryoCodex also achieved the highest mean sequence recall of 31.7%, compared with 25.8% for EMReady2 and 15.5% for the deposited maps. Because sequence recall imposes the additional requirement of correct residue identity on spatially matched residues, the higher sequence recall further indicates that CryoCodex improves local density features sufficiently to support more accurate residue identification, beyond simply increasing the number of spatially traceable residues. Taken together, these results demonstrate that the benefits of CryoCodex extend beyond improvements in map-level quality metrics: the enhanced maps contain structural features that are substantially more informative for independent automated model building, enabling both more complete structural tracing and more accurate residue assignment across the benchmark. Mean values for all methods are provided in Supplementary Table 3, and complete per-chain results are provided in Supplementary Data 3.

**Fig. 4.**
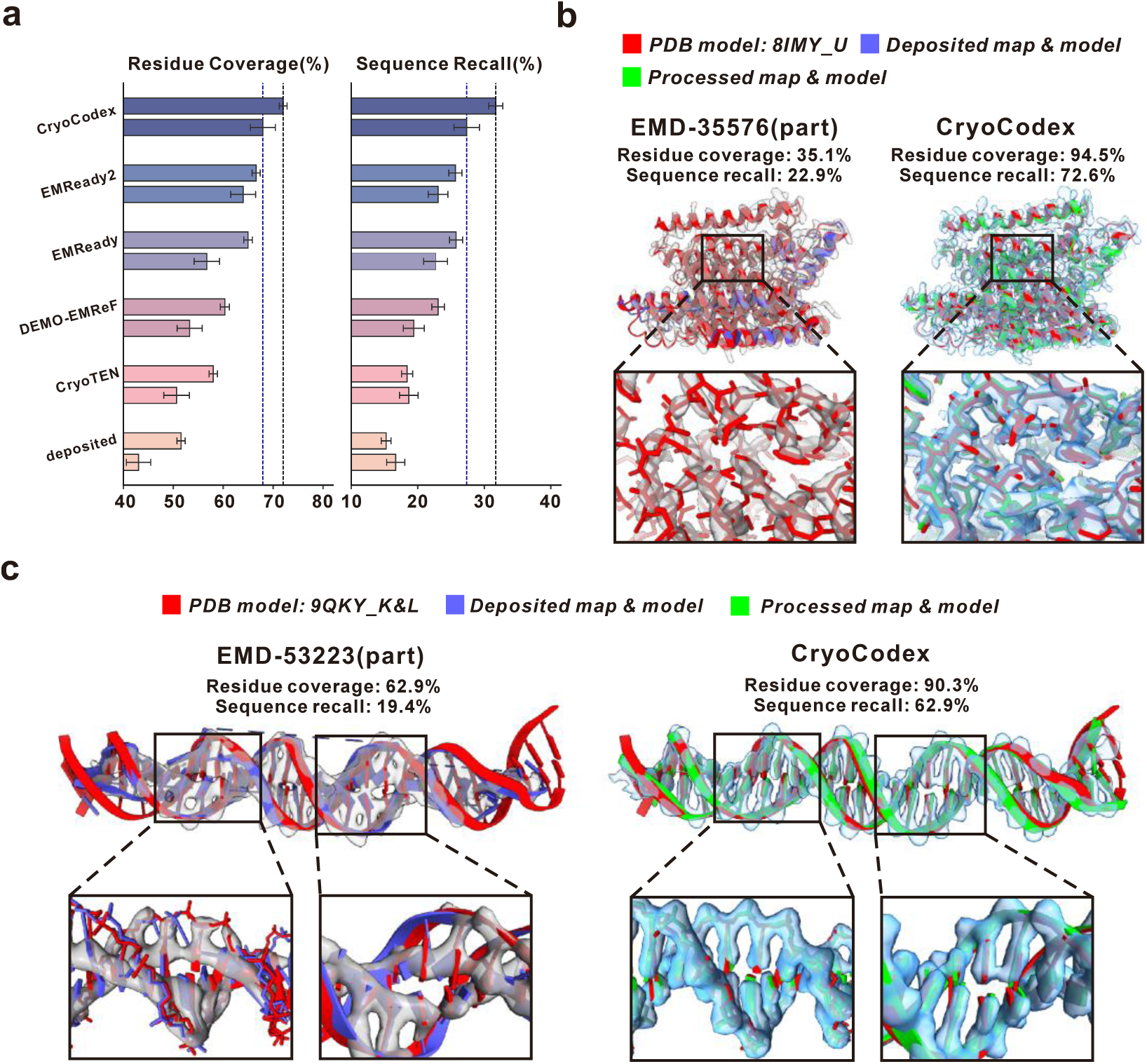
| Improvement in map interpretability. **a** Residue coverage and sequence recall across different methods. For each method, the upper and lower bars represent the common overall subset (n = 761) and nucleic-acid subset (n = 102), respectively. Dashed lines indicate the mean CryoCodex performance on the corresponding datasets; error bars show SEM. **b** Protein example EMD-35576 (PDB 8IMY, chain U), comparing the deposited and CryoCodex-processed maps and the corresponding Phenix-built models with the reference structure. **c** DNA example EMD-53223 (PDB 9QKY, chains K and L), shown in the same manner, with enlarged views highlighting local differences in model building.

Fig. 4b illustrates a representative example for EMD-35576, a 3.22 Å cryo-EM map of the *substrate-bound human glycosylphosphatidylinositol transamidase complex*, focusing on its PIGU subunit (PDB ID: 8IMY, chain U). The atomic structure was determined using template fitting rather than *de novo* modeling, likely due to the challenges illustrated below. Automated *de novo* model building from the deposited map recovered only a limited fraction of the reference structure, yielding a residue coverage of 35.1% and a sequence recall of 22.9%. Several α-helical segments remained incompletely traced in the resulting model. In contrast, automated modeling from the CryoCodex-processed map recovered substantially more of the reference structure, increasing residue coverage to 94.5% and sequence recall to 72.6%. The enlarged views of the boxed region in Fig. 4b further illustrate the local density changes underlying this improvement. In the deposited map, density along parts of the backbone is weak or locally discontinuous, while several residue-associated density features are poorly defined. After CryoCodex processing, the density becomes more continuous along the reference backbone, and local residue-associated features become more clearly defined. These improvements provide stronger spatial constraints for backbone tracing and residue assignment, thereby enabling substantially more complete and accurate *de novo* model building.

### CryoCodex enhances nucleic acid density and downstream model building

Nucleic acid density can be particularly challenging^7, 30–35^ to interpret because the four canonical bases provide less chemical diversity than amino acids, while backbone and base features become increasingly difficult to distinguish at moderate-to-low resolution. We therefore separately evaluated CryoCodex on a dedicated subset of 22 test maps in which nucleic acid residues accounted for more than 10% of the corresponding atomic model, with reported resolutions ranging from 3.08 to 7.50 Å.

As shown in Fig. 2a,d,e and Supplementary Table 2, CryoCodex consistently achieved the best performance across the evaluated metrics on this nucleic-acid-containing subset. For FSC-0.5, comparisons were performed on the common subset of 20 maps for which valid results were available for all evaluated methods. CryoCodex achieved the lowest mean FSC-0.5 resolution of 3.43 Å, compared with 4.02 Å for the second-best method, EMReady2, and 5.45 Å for the deposited maps. CryoCodex improved the FSC-0.5 resolution by more than 1 Å relative to the deposited maps in 80.00% of cases, compared with 35.00% for EMReady2. Q-score statistics were calculated on all 22 maps. CryoCodex achieved the highest mean Q-score of 0.522, compared with 0.474 for EMReady2 and 0.427 for the deposited maps. In addition, 86.36% of CryoCodex-processed maps showed a Q-score improvement greater than 0.05, and 68.18% showed an improvement greater than 0.08. Complete per-target FSC-0.5 and Q-score results for the nucleic-acid-containing subset are provided in Supplementary Data 2.

At the chain level, we further evaluated automated *de novo* model building on the 102 nucleic acid chains in the common modeling subset described above. Models built from the CryoCodex-enhanced maps achieved a mean residue coverage of 69.1% and a mean sequence recall of 27.8%, compared with 64.7% and 23.3% for EMReady2, and 43.8% and 17.0% for the deposited maps, respectively (Fig. 4a and Supplementary Table 3). These improvements indicate that the enhanced nucleic acid density supports both more complete structural tracing and more accurate nucleotide assignment, providing downstream evidence that the improvements observed in map-level metrics translate into enhanced structural interpretability for nucleic acid regions.

Fig. 3d–f illustrate a representative protein–DNA example from the *Drosophila melanogaster* P-element transposase strand transfer complex (EMD-20321, PDB ID: 6PE2). At a lower contour level (Fig. 3d), the deposited map contains substantial diffuse and peripheral density around the complex, whereas the CryoCodex-processed map exhibits a cleaner and more compact molecular envelope with clearer separation from the surrounding density. Differences in the DNA region become more evident at the higher contour level and in the enlarged views shown in Fig. 3e. In the deposited map, the density surrounding the phosphodiester backbone is relatively broad, and backbone- and base-associated densities are locally merged, making the trajectory of the DNA backbone and the spacing between adjacent bases difficult to distinguish. In contrast, the CryoCodex-processed map shows a more continuous and better-defined phosphodiester backbone, with density more closely following the reference DNA model. The enlarged view further reveals clearer separation of individual base-associated features, making the periodic spacing and stacked arrangement of adjacent bases more discernible. Consistent with these visual observations, the unmasked map– model FSC curves show increased agreement with the reference structure over a broad range of spatial frequencies after CryoCodex processing (Fig. 3f). The FSC-0.5 resolution improved from 4.43 Å to 3.49 Å, while the Q-score increased from 0.411 to 0.599.

Fig. 4c further illustrates how improvements in nucleic acid density facilitate downstream automated model building, using EMD-53223 (PDB ID: 9QKY, chains K and L), derived from a *DNA-bound Nuclear Factor 1-X (NFI-X) complex*. *De novo* modeling from the deposited map yielded a residue coverage of 62.9% and a sequence recall of 19.4%, whereas modeling from the CryoCodex-processed map increased these values to 90.3% and 62.9%, respectively. The enlarged regions at the bottom of Fig. 4c provide a direct explanation for these improvements. In one representative region, the deposited map contains weak and locally discontinuous density along the phosphodiester backbone, resulting in incomplete or deviated backbone tracing in the Phenix-built model. After CryoCodex processing, continuous density is restored along the corresponding backbone path, allowing the automatically built model to follow the reference DNA more closely. In another enlarged region, the deposited map provides poorly defined density around an individual base, as reflected by a mean deposited Q-score of only 0.487 in the corresponding region, resulting in an inaccurate base placement or orientation relative to the reference model. The CryoCodex-enhanced density displays a more clearly defined base-associated feature, with the mean Q-score in the corresponding region increased to 0.644 and the spatial orientation more clearly resolved, thereby providing stronger local constraints for nucleotide placement. Thus, the improvement is not limited to recovering a more continuous DNA backbone, but also extends to local base features required for more accurate nucleotide identification, consistent with the pronounced increase in sequence recall.

Taken together, these examples show that CryoCodex improves nucleic acid density at multiple structural levels, from the overall molecular envelope and continuity of the phosphodiester backbone to the separation and definition of individual base-associated features. These improvements provide more informative density for both visual interpretation and automated model building.

### CryoCodex enhances weak structural features without noise overfitting or structural hallucination

Deep-learning-based map enhancement can improve the visibility of weak structural features, but two related concerns remain. First, a highly flexible model may overfit unstable or half-map-specific density fluctuations and amplify them as apparently meaningful structural features. Second, learned structural priors may lead to structural hallucination, in which ambiguous or poorly supported density is artificially regularized into plausible but unsupported macromolecular features. We therefore evaluated these two risks separately. Half-map cross-validation was used to assess whether CryoCodex shows a systematic tendency to fit half-map-specific noise, whereas a representative weak-signal region was examined to evaluate whether the model forces poorly supported density into canonical structural patterns.

Half-map data were available for 90 maps in the test set. For each dataset, the two independently reconstructed unprocessed half-maps were denoted U-1 and U-2, and CryoCodex was independently applied to each half-map to yield P-1 and P-2, respectively. Unmasked map-map FSC curves were then calculated for four pairwise combinations. U-1 versus U-2 represents the baseline correlation between the two independent experimental half-maps. The cross-pair comparisons P-1 versus U-2 and U-1 versus P-2 assess whether features enhanced from one half-map remain correlated with signals present in the independently reconstructed, unprocessed counterpart. P-1 versus P-2 evaluates the reproducibility of the enhancement across the two independent inputs. Because both processed half-maps are generated by the same trained network, P-1/P-2 agreement is interpreted together with the two processed-unprocessed cross comparisons when assessing enhancement reliability.

In this analysis, FSC is used primarily as a measure of Fourier-space correlation. We restricted the comparative analysis to the common subset of 76 maps for which all four FSC calculations yielded valid results. The statistical summaries and distributions on these maps are shown in Fig. 5. Complete per-target results for all available half-map datasets are provided in Supplementary Data 4. The FSC-0.5 crossing positions shown in Fig. 5 are therefore treated as descriptive summaries of relative shifts in the correlation curves rather than as direct estimates of physical resolution. In particular, crossing positions extending beyond the conventional isotropic Nyquist limit are not interpreted as physical resolution values. Across the 76 datasets, the processed-unprocessed cross comparisons retained substantial Fourier-space correlation with the independent unprocessed half-maps, whereas P-1 and P-2 showed still stronger mutual agreement (Fig. 5a). This combination of cross-pair consistency and processed– processed reproducibility indicates that CryoCodex enhancement remains anchored to information present in independent experimental observations while producing highly reproducible structural features.

**Fig. 5.**
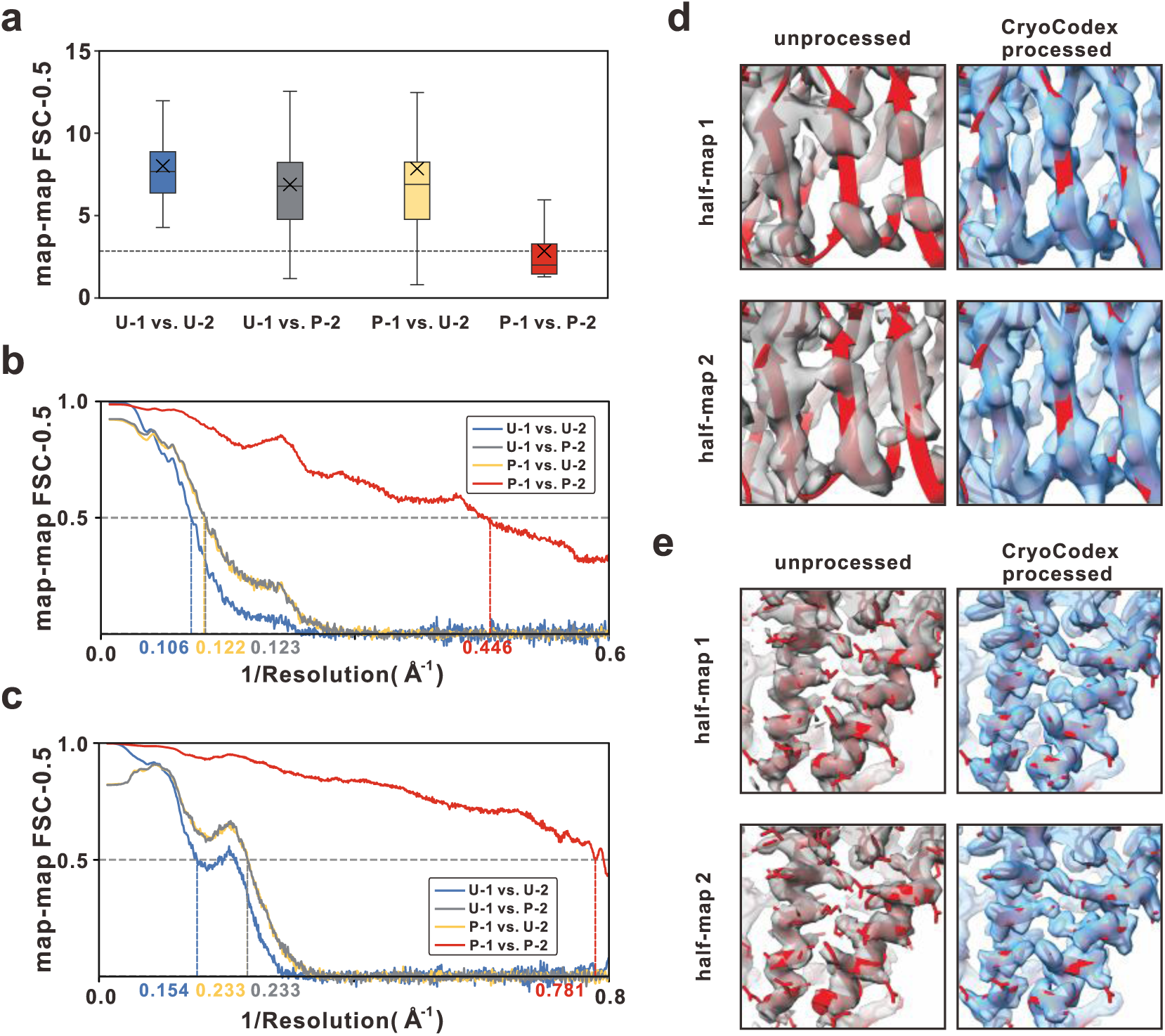
| Half-map cross-validation of enhancement reproducibility and consistency. **a** Box-and-whisker plot showing the distribution of FSC-0.5 cutoffs from unmasked map–map FSC curves across the common subset of 76 half-map datasets for which valid FSC-0.5 crossings were available for all four pairings. Boxes indicate the interquartile range, center lines indicate the median, whiskers extend to the most extreme data points within 1.5 times the interquartile range, and × symbols indicate the mean. U-1/U-2 represent the original unprocessed half-maps, and P-1/P-2 represent the CryoCodex-enhanced half-maps. **b, c** Map–map FSC curves of the four half-map combinations for EMD-14800 (**b**) and EMD-16429 (**c**), with FSC-0.5 spatial frequencies indicated below the dashed lines. **d, e** Detailed density visualizations of unprocessed and CryoCodex-enhanced half-maps 1 and 2, superimposed on the corresponding deposited models for EMD-14800

The half-map design also provides a direct way to examine noise overfitting. Features arising predominantly from random fluctuations specific to U-1 would not be expected to retain consistent correlation with the independently reconstructed U-2 after processing, and the same reasoning applies symmetrically to features originating from U-2. The persistence of correlation in both processed-unprocessed cross pairs therefore argues against systematic fitting of half-map-specific random noise as the dominant source of the observed enhancement. At the same time, the stronger agreement between P-1 and P-2 indicates that CryoCodex consistently emphasizes structural information shared by the two independent observations. Thus, although the half-map analysis cannot establish the correctness of every individual enhanced feature, it provides evidence that the overall improvement is not primarily driven by fitting noise unique to a single half-map.

Fig. 5b,d illustrate this behavior for EMD-14800, a 4.32 Å cryo-EM map of the *bacteriophage* T5 tail-tip region with its central fiber bent following interaction with the bacterial receptor *FhuA*. In the two unprocessed half-maps, density in the highlighted β-sheet-rich region is weak and locally discontinuous, making the backbone trajectory difficult to follow consistently. After independent CryoCodex processing, both half-maps exhibit more continuous density along the corresponding β-sheet region, with closely matching backbone trajectories around the reference atomic model (Fig. 5d). The P-1/P-2 FSC curve is correspondingly elevated relative to the U-1/U-2 baseline over a broad range of spatial frequencies (Fig. 5b). Importantly, the processed-unprocessed cross-pair correlations remain comparable to, and in some spatial-frequency ranges even exceed, the correlation between the two independently reconstructed unprocessed half-maps. This further supports that the enhanced β-sheet features reflect structural information already present in the original data, rather than simply arising from similarity between the two processed outputs.

EMD-16429, a 3.36 Å cryo-EM map of the *Hantaan* virus RNA-dependent RNA polymerase in a replication pre-initiation state, provides a second representative example involving α-helical and side-chain-associated density (Fig. 5c, e). Although the α-helical backbone is discernible in both unprocessed half-maps, many local side-chain-associated features are weak or poorly defined. Following independent CryoCodex processing, both half-maps show more clearly defined density at corresponding side-chain positions and closely matching local orientations around the reference model (Fig. 5e). Consistently, the processed half-maps exhibit stronger mutual FSC over much of the sampled spatial-frequency range, while the processed-unprocessed cross comparisons continue to show agreement with the independent unprocessed observations (Fig. 5c). The reproducible appearance of these local features in both half-maps, together with their cross-half-map support, is consistent with enhancement of weak structural information already present in the experimental data rather than arbitrary amplification of half-map-specific fluctuations.

Beyond noise overfitting, a related concern for supervised enhancement models is structural hallucination in regions where the experimental density is ambiguous or weakly supported. To examine this behavior, we analyzed EMD-19811 (PDB ID: 8S8P), a 3.11 Å cryo-EM map of the *budding yeast Ku-Rap1-DNA complex involved in telomere protection*, focusing on a challenging peripheral region containing a helical segment and an adjacent flexible loop (Supplementary Fig. 2). At a lower contour level, CryoCodex substantially suppresses diffuse background density while preserving the overall molecular envelope. More importantly, the enlarged views show that CryoCodex enhances the contrast of the peripheral region and clarifies its local density boundaries without forcing the ambiguous signal into an artificially regularized continuous helix or other canonical structural features. At a higher contour level, structurally well-supported regions become more sharply defined, while the peripheral density retains the heterogeneous characteristics present in the experimental map. This selective enhancement behavior shows that CryoCodex improves structural visibility without indiscriminately regularizing ambiguous density toward idealized structural patterns.

Taken together, the half-map cross-validation and weak-signal case provide complementary evidence for the structural fidelity of CryoCodex enhancement. The reproducibility observed across independently processed half-maps, together with the retained correlation between each processed half-map and the opposite unprocessed half-map, indicates that the enhancement is unlikely to be dominated by fitting half-map-specific random noise. Meanwhile, the weak-signal example shows that CryoCodex enhances structurally supported density without indiscriminately imposing idealized structural patterns in ambiguous regions. These observations support the robustness of CryoCodex in enhancing weak structural information while preserving consistency with the underlying experimental data.

### CryoCodex predicts local density quality and molecular regions alongside map enhancement

Global map-level metrics provide useful overall summaries of cryo-EM map quality but do not capture the substantial spatial heterogeneity that can occur within individual maps. Such spatial variation can persist after enhancement because different regions may inherently differ in signal strength and structural flexibility, leading to differences in local interpretability. To characterize these spatial variations, CryoCodex predicts two voxel-wise quality maps: the local quality of the input density map (in-score) and that of the enhanced density map (out-score). In parallel, the network predicts a molecular mask to delineate molecular regions from the surrounding background. These two types of prediction serve distinct purposes: the in-score and out-score provide spatially resolved estimates of density quality before and after enhancement, whereas the molecular mask identifies the spatial extent of molecular density. The quality branches are supervised using voxel-wise 3D structural-similarity targets derived from the experimental and PDB-derived simulated density maps, while the molecular mask is supervised using PDB-derived molecular regions (Methods).

We first evaluated whether the in-score captures local quality variations in the deposited maps. CryoCodex was compared with CryoRes^24^, an end-to-end local resolution estimation method, on all 126 test maps. For each map, voxel-wise predictions were mapped to the corresponding reference atomic coordinates using nearest-neighbor sampling, and their Spearman correlations with per-atom Q-scores were calculated. Because lower CryoRes values indicate better local resolution, the sign of its Spearman coefficient was reversed so that larger positive values consistently indicate stronger correspondence with Q-score. As shown in Fig. 6a, CryoCodex achieved a mean per-map Spearman correlation of 0.22, compared with 0.18 for CryoRes. When all atoms from the test set were pooled, the corresponding correlations were 0.48 and 0.43, respectively. These results indicate that the in-score captures spatial variation in local map quality and shows stronger correspondence with atom-level Q-score than CryoRes.

**Fig. 6.**
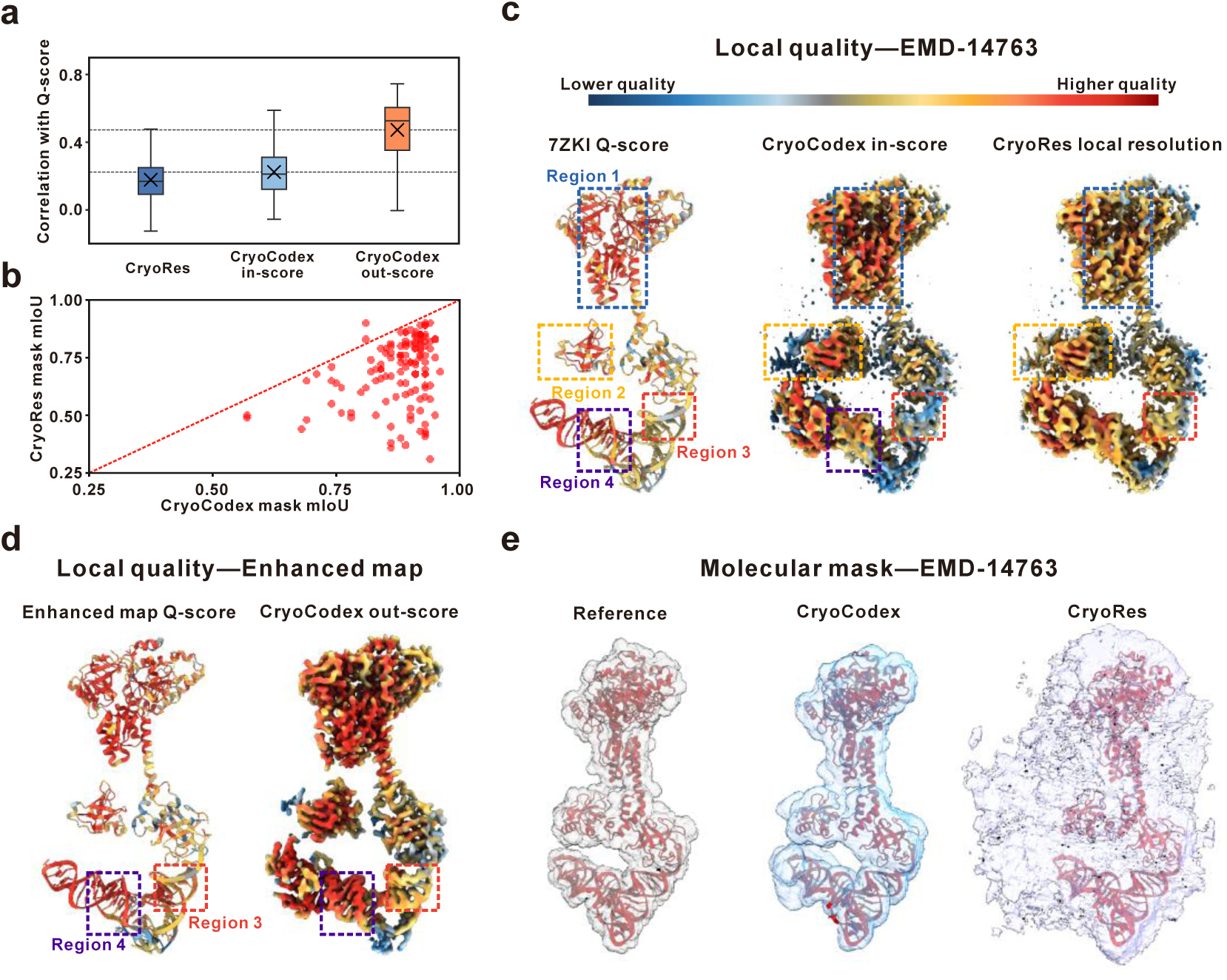
| Evaluation of local map quality estimation and molecular mask prediction. **a** Spearman correlation coefficient between predicted local quality metrics and per-atom Q-scores across the test set (CryoRes signs are inverted). Boxes indicate the interquartile range, center lines indicate the median, whiskers extend to the most extreme data points within 1.5 times the interquartile range, and × symbols indicate the mean. **b** Molecular mask prediction accuracy (mIoU) comparison between CryoCodex and CryoRes. **c** Local quality estimation on the deposited map of EMD-14763. Left to right: atomic model colored by Q-score, and maps colored by CryoCodex In-score and CryoRes local resolution. **d** Local quality assessment on the enhanced map of EMD-14763 (left: model colored by Q-score; right: map colored by CryoCodex Out-score). Regions 3 and 4 highlight the local quality improvement compared to (**c**). **e** Molecular masks of EMD-14763: Reference mask, CryoCodex, and CryoRes.

We next evaluated whether the out-score provides corresponding quality information for the enhanced density maps. Using per-atom Q-scores calculated from the CryoCodex-enhanced maps as the reference, the out-score achieved a mean per-map Spearman correlation of 0.47 and a pooled atom-level correlation of 0.71 (Fig. 6a). These correlations indicate that the out-score captures spatial variation in quality within the enhanced maps. The out-score therefore provides information complementary to the enhanced density itself, allowing regions of relatively high or low predicted quality to be identified directly within the processed map.

The molecular mask was evaluated separately as a molecular-region prediction task. Across the test set, CryoCodex achieved a mean intersection-over-union (mIoU) of 0.88 and a foreground IoU of 0.79, compared with 0.69 and 0.54 for CryoRes, respectively (mIoU shown in Fig. 6b). The per-map comparison further shows that this advantage is broadly distributed across the benchmark, with most samples lying below the equality line and therefore exhibiting higher mIoU for CryoCodex. These results indicate that CryoCodex consistently delineates molecular regions across diverse experimental maps while excluding surrounding non-molecular density. Detailed per-target results for local quality prediction and molecular mask prediction are provided in Supplementary Data 5 and 6, respectively.

EMD-14763, a 3.6 Å cryo-EM map of a translation initiation subcomplex comprising the archaeal factors aIF1A and aIF5B bound to initiator tRNA, provides a representative example of the spatial information captured by these predictions (Fig. 6c–e). On the deposited map, the CryoCodex in-score achieved a Spearman correlation of 0.40 with the corresponding per-atom Q-scores, compared with 0.28 for CryoRes (Fig. 6c). Beyond the overall correlation, the predicted in-score captured several local quality patterns reflected by the per-atom Q-scores. Region 1, which exhibits relatively high Q-scores, was assigned correspondingly high in-score values with clear spatial separation from the surrounding lower-quality regions. In Region 2, the in-score reproduced the gradual spatial transition in local quality while assigning low scores to adjacent diffuse density. In the nucleic-acid-associated Region 3, where the deposited map exhibits locally reduced Q-scores, CryoCodex similarly identified a localized low-quality region, whereas this variation was less clearly distinguished by CryoRes.

Comparison of the deposited and enhanced maps further illustrates the complementary roles of the in-score and out-score. For the enhanced map, the out-score showed a Spearman correlation of 0.63 with the corresponding per-atom Q-scores (Fig. 6d). In Regions 3 and 4, local increases in Q-score after enhancement were accompanied by corresponding increases from in-score to out-score. The spatial agreement between these changes indicates that the paired quality predictions can reveal where local map quality has improved following enhancement, rather than providing only a global assessment of the processed map.

The molecular mask prediction for the same example provides an independent assessment of spatial boundary delineation (Fig. 6e). CryoCodex achieved foreground and background IoUs of 0.78 and 0.98, respectively, compared with 0.19 and 0.61 for CryoRes. Consistent with these values, the CryoCodex mask closely followed the reference molecular envelope while excluding much of the surrounding non-molecular density, whereas the CryoRes prediction showed substantially more false-positive mask voxels outside the reference molecular boundary.

Together, these results show that CryoCodex couples density enhancement with spatially resolved quality assessment while independently identifying molecular regions. The in-score and out-score capture local quality heterogeneity before and after enhancement, whereas the molecular mask delineates the spatial extent of molecular density. This unified framework therefore provides both the enhanced density and complementary spatial information for interpreting its local quality.

### Ablation experiments show MMVQ reduces overfitting and improves background suppression

To assess the contribution of MMVQ, we constructed an ablation model by removing the MMVQ pathway and its associated gated fusion. We compared the training dynamics of the full and ablated models using the two loss terms associated with the density enhancement task (Supplementary Fig. 3a–d). During training, the model without MMVQ reached lower Smooth *L*1 and SSIM losses for density enhancement than the full model, whereas the corresponding validation losses remained comparatively higher. This pattern—lower training losses but poorer validation performance—suggests a stronger tendency toward overfitting in the absence of MMVQ. This behavior is consistent with the structured discrete constraints introduced by MMVQ. By quantizing latent representations across multiple feature subspaces and spatial scales, MMVQ limits unconstrained adaptation of the continuous latent representation to sample-specific fluctuations, thereby promoting representations that generalize more effectively beyond the training data.

Supplementary Fig. 3e provides a representative example consistent with this interpretation. The deposited 4.1 Å cryo-EM map of *retinal-free channelrhodopsin ChRmine reconstituted in a lipid nanodisc* (EMD-25135) contains substantial diffuse density surrounding the molecular region. The ablated model without MMVQ retains pronounced residual background density in the highlighted region, whereas the full CryoCodex model more effectively suppresses this surrounding density while preserving the molecular density. Together with the training and validation behavior, this example supports the role of MMVQ in constraining the enhancement process and reducing the tendency of the network to fit unstable background fluctuations.

### Computational efficiency

To evaluate computational efficiency under a practical and consistent memory constraint, we compared the running time of CryoCodex with four representative deep learning-based post-processing methods—CryoTEN, DEMO-EMReF, EMReady, and EMReady2—across all 126 test maps (Supplementary Fig. 4). Because these methods differ substantially in their input box sizes, sliding strides, and memory requirements, allowing each method to use unrestricted GPU memory would make the comparison strongly dependent on hardware capacity and achievable batch size. We therefore constrained GPU memory usage to no more than 8 GB for all methods to better reflect a practical inference scenario with limited computational resources. All running-time measurements were performed on the same NVIDIA A800 GPU. Each method retained its default inference configuration, while the input batch size was independently set to the maximum value supported within the 8 GB memory limit. Under this setting, the maximum batch sizes for CryoTEN, DEMO-EMReF, EMReady, EMReady2, and CryoCodex were 111, 6, 45, 28, and 9, respectively.

CryoTEN showed the shortest mean running time at 44.80 s per map, followed by CryoCodex at 74.87 s, EMReady2 at 106.69 s, EMReady at 173.96 s, and DEMO-EMReF at 191.30 s (Supplementary Fig. 4). Complete per-target running times for all evaluated methods are provided in Supplementary Data 7. CryoCodex achieved the second-shortest running time among the evaluated methods and was faster than DEMO-EMReF, EMReady, and EMReady2. Notably, this computational efficiency was achieved while CryoCodex also showed the best overall map-enhancement performance among the compared methods in the preceding evaluations. Moreover, a single CryoCodex run simultaneously generates the enhanced density map, in-score, out-score, and molecular mask, rather than producing only an enhanced map. Taken together, these results show that CryoCodex maintains competitive running time despite performing multiple prediction tasks and achieving superior enhancement performance.

## Discussion

In this work, we presented CryoCodex, a deep-learning framework that unifies cryo-EM map enhancement, local quality estimation, and molecular masking within a single network. At the core of the framework is the multi-head, multi-scale vector-quantization (MMVQ) module, which represents latent features through a learned codebook across feature subspaces and spatial scales. By combining coarse-to-fine residual quantization with gated fusion, CryoCodex imposes discrete constraints on the learned representation while preserving the continuous information needed to capture fine structural detail. This design directly addresses a central challenge in cryo-EM map enhancement: reducing the tendency to fit unstable density fluctuations without over-suppressing weak yet genuine structural features. We assessed CryoCodex on a held-out benchmark of 126 experimental maps and found that it achieved superior performance across all considered tasks.

Looking forward, several directions remain open. Extending the framework to handle larger and more heterogeneous datasets, incorporating additional validation metrics, and integrating downstream model-building pipelines could further improve its practical utility. More broadly, we expect that coupling enhancement with interpretable, region-specific quality estimation will become increasingly important as cryo-EM continues to push toward the analysis of highly flexible and heterogeneous macromolecular assemblies.

## Methods

### Network architecture

Figure 1a illustrates the overall architecture of CryoCodex. MMVQ-UNet is a vector-quantized U-Net that maps continuous feature representations onto a finite, discrete codebook. This vector-quantization strategy is motivated by the assumption that continuous features in cryo-EM density maps can be decomposed into a set of recurring structural patterns with superimposed stochastic noise^25^; vector quantization therefore provides an effective means of extracting more representative prototype features while suppressing noise. The U-Net backbone is adapted from the RU-Net originally developed in CryoAtom2 ^7^ for key-atom localization. For computational feasibility, we partition the density map into multiple input density boxes of size 48 × 48 × 48. The encoder, composed of convolutional layers, first extracts continuous latent features, which are then processed by a multi-head, multi-scale vector-quantization (MMVQ) module to derive representative prototype features from the learned codebook. The quantized and continuous representations are adaptively fused through a gated fusion block, followed by a global attention module that further updates the feature representations. Finally, the decoder restores the spatial resolution and uses four task-specific output heads to predict the enhanced density box, input-map local quality, enhanced-map local quality, and molecular region mask, respectively.

### MMVQ module

The multi-scale quantization scheme was originally proposed by VAR^36^ in the field of image generation. In CryoCodex, we extend this strategy to three-dimensional latent representations and combine it with multi-head feature partitioning. Firstly, the encoder produces a continuous latent representation after three downsampling stages, with the following shape: *E* ∈ ℝ*^B^*^×*C*×6×6×6^. Here, *B* denotes the batch size and *C* denotes the number of feature channels. Next, inspired by the multi-head design, the latent representation is evenly divided into ℎ = 8 groups along the channel dimension. Each feature group therefore contains *d* = *C*/ℎ channels and is processed by an independent quantization head. Finally, we describe the multi-scale residual quantization strategy in detail, which provides finer-grained feature representations than conventional quantization methods. At the multi-head level, multi-scale residual quantization is performed independently within each head, while a single codebook C is shared across all quantization heads and spatial scales.

Taking a single quantization head as an example, quantization is performed recursively from coarse to fine over six spatial scales, indexed by *k* = 1, …, 6 (Fig. 1b). For clarity, the quantization-head index is omitted in the following formulation. The residual before the first quantization stage is initialized as the corresponding encoder feature, denoted by *R*_0_. At each scale *k*, the residual *R_k_*_−1_ from the preceding scale is first resampled to the current *k* × *k* × *k* grid. Each spatial feature vector is then assigned to its nearest vector in the shared codebook. The resulting discrete representation is restored to the original latent resolution and subtracted from the current residual. The complete recursive procedure is expressed as:

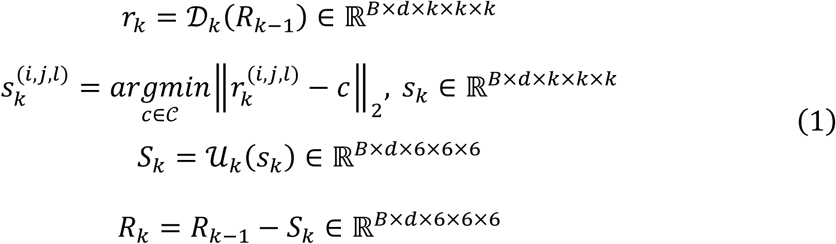

Here, *D_k_*(⋅) denotes spatial downsampling from the original 6 × 6 × 6 latent grid to the current *k* × *k* × *k* grid, and *U_k_*(⋅) upsamples the quantized representation to the original latent resolution using trilinear interpolation. (*i*, *j*, *l*) denotes a spatial location on the current-scale grid. The shared codebook C contains ∣ C ∣= 4096 updatable feature vectors, each with dimension *d*. At every spatial position, the nearest-neighbor assignment in Eq. (1) selects the codebook vector with the minimum Euclidean distance to the corresponding continuous feature vector, effectively replacing it with a representative discrete feature that suppresses noise.

The recursive update of *R_k_* enables successive scales to operate on information that has not been represented at preceding coarser scales. Specifically, *R_k_*_−1_ is resampled and quantized at scale *k*, the resulting quantized component *S_k_* is removed, and the remaining residual *R_k_* is passed directly to scale *k* + 1. The spatial granularity therefore progressively increases from 1^3^ to 6^3^, allowing the remaining residual information to be represented from coarse to fine.

The quantized components obtained across all spatial scales are accumulated to produce the output of the corresponding quantization head:

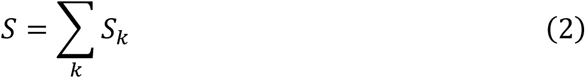

The outputs of all quantization heads are subsequently concatenated along the channel dimension. The resulting quantized representation is then combined with the corresponding continuous encoder features through the gated fusion block. Specifically, the quantized features are used to generate a gating weight through a grouped 1 × 1 × 1 convolution followed by a sigmoid activation, while the continuous features are transformed by a 3 × 3 × 3 convolution. The gated continuous features are then added to the quantized representation before being passed to the subsequent network layers. The gating convolution uses eight groups, corresponding to the number of quantization heads. During backpropagation, gradients through the discrete quantization operation were propagated to the encoder using a straight-through estimator (STE). The codebook was shared across all quantization heads and spatial scales and was updated exclusively using exponential moving averages (EMA). No additional codebook or commitment loss was used.

### Training and benchmark datasets

The data used in this study were obtained from the Electron Microscopy Data Bank (EMDB)^1^ and the Protein Data Bank (PDB)^29^. Cryo-EM density maps with reported resolutions ranging from 3 to 8 Å were initially collected and filtered according to the following criteria: (1) the reported resolution was determined using the FSC-0.143 criterion; (2) each density map had a uniquely corresponding PDB atomic model; (3) the atomic model contained complete structural information without unknown atoms, residues, or missing chains; (4) the density map had orthogonal coordinate axes; (5) the CC_masks^26^ between the experimental density map and the corresponding atomic model was greater than 0.5; (6) the map-model correlation calculated using Chimera^28^ was greater than 0.6; and (7) the total number of Cα and C4′ atoms in the structure did not exceed 10,000.

To reduce sequence redundancy, protein and nucleic acid chains were clustered separately using MMseqs2^37^. For protein chains, sequence coverage and sequence identity thresholds were set to 80% and 30%, respectively, whereas both thresholds were set to 80% for nucleic acid chains. Chains assigned to the same cluster were considered sequence-redundant, and two map–structure pairs were considered redundant if at least one pair of chains between them satisfied the corresponding redundancy criterion. During overall dataset curation, a greedy selection procedure was applied to remove redundant structures, with priority given to structures containing nucleic acids and, within the same category, to lower-resolution maps. After filtering and redundancy removal, a total of 1658 non-redundant cryo-EM density map– structure model pairs were retained.

From this dataset, 126 samples were held out as the independent test set. Approximately 48% of the test maps had reported resolutions worse than 4 Å, approximately 21% had resolutions worse than 5 Å, and approximately 17% corresponded to structures in which nucleic acid residues accounted for more than 10% of the atomic model. Of the remaining samples, 66 were assigned to the validation set and the others were used for model training. For downstream automated model-building evaluation, all structural chains in the 126-map test set were extracted, yielding 771 chains, including 668 protein chains and 103 nucleic acid chains. For each chain, a chain-specific local density map was extracted from the original density map using the corresponding atomic coordinates, retaining the density region within 4 Å of any atom in the chain. These chain-specific density maps were used for the map-interpretability evaluation.

No additional restriction was imposed on the experimental reconstruction method during dataset selection. Consequently, the dataset was not limited to single-particle analysis (SPA); however, maps obtained using other reconstruction approaches, including subtomogram averaging and helical reconstruction, represented only a small fraction of the final dataset after filtering. We therefore did not perform separate evaluations according to reconstruction method.

### Training and Inference

All raw cryo-EM density maps were resampled to a uniform voxel size of 1 Å using trilinear interpolation. Negative voxel values were truncated to zero, and each map was normalized by its 99.999th percentile. For training and validation sample preparation, the minimum bounding cube enclosing the corresponding PDB structure was first determined on the resampled density map grid and expanded outward by 15 Å along each dimension. The corresponding region was then cropped to exclude extensive background areas that provide no effective training signal. A 70 × 70 × 70 sliding window with a stride of 35 was subsequently used to extract candidate sub-volumes, and only sub-volumes with a maximum density value greater than zero were retained. During training, 48 × 48 × 48 boxes were randomly cropped from these sub-volumes and augmented with random rotations in 90° increments, whereas center cropping was used during validation. During inference, approximate molecular regions can be estimated to exclude a large number of background boxes and accelerate inference, inspired by the ModelAngelo auto-mask procedure ^8^. The cropped map was then processed using 48 × 48 × 48 boxes with a sliding stride of 12, which were used as the inference inputs. For convenience, the input density box is hereafter denoted as *B_d_*, and the output density box as *B_o_*. The training labels and loss functions are described below.

#### Training labels

The supervision label corresponding to the input density map (*B_d_*) is denoted as *B_s_*. It is derived from a simulated density map generated from the atomic coordinates corresponding to the same spatial region of the input density map ^11^. The binary molecular mask, denoted as *B_m_*, was constructed by assigning a value of 1 to voxels within 4 Å of any atom and 0 to the remaining background voxels. A reweighted mask 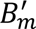, assigning values of 1 and 0.001 to foreground and background voxels, respectively, was further used for the density enhancement loss.

For the input density box (*B_d_*), its local quality is defined using the Structural Similarity Index Measure (SSIM) and is denoted as *Q_d_*_,*s*_, which is specifically defined as follows:

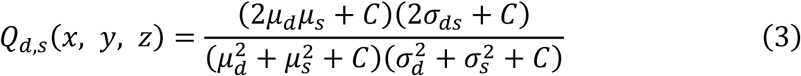

where *C* = 1 × 10^−6^ is a constant used for numerical stability. To compute the local quality, a Gaussian kernel (5 × 5 × 5) was applied to both *B_d_* and *B_s_* to obtain voxel-wise local statistics. For each spatial coordinate (*x*, *y*, *z*), the local means *μ_d_* and *μ_s_*, local variances 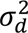 and 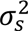, and covariance *σ_ds_* were calculated. Similarly, for the output density box (*B_o_*), the corresponding local quality *Q_o_*_,*s*_ can be defined in a manner analogous to Eq. (3).

#### Loss functions

For the density enhancement task, a combination of weighted Smooth *L*_1_ loss and weighted SSIM loss was used for optimization. The weighted Smooth *L*_1_ loss is defined as

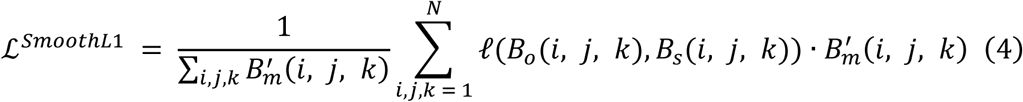

where *N* = 48, and *l*(*B_o_*(*i*, *j*, *k*), *B_s_*(*i*, *j*, *k*)) denotes the Smooth *L*_1_ distance between density values at position (*i*, *j*, *k*):

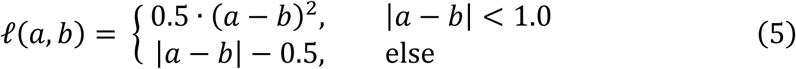

The weighted SSIM loss for the density enhancement task is defined as:

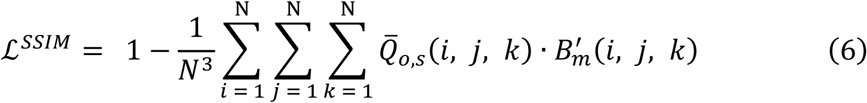

where *Q̅_o_*_,*s*_ is calculated analogous to *Q_o_*_,*s*_, except that the mean terms are omitted and an 11 × 11 × 11 convolution window is used to capture broader contextual information:

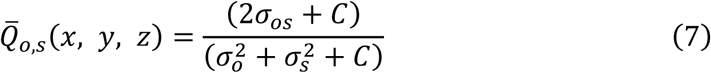

where *C* is the same numerical stability constant as in Eq. (3).

The input- and enhanced-map local quality prediction tasks were each optimized using a Smooth *L*_1_ loss, whereas the molecular mask prediction task was optimized using binary cross-entropy (BCE) loss. The overall multitask objective was defined as the unweighted sum of the five loss terms. The model was trained on six NVIDIA A800 GPUs using the Adam optimizer with an initial learning rate of 5 × 10^−4^. Gradient accumulation was used, resulting in an effective batch size of 264. Training was terminated when any of the following conditions was met: the learning rate decreased to 1 × 10^−5^, the maximum number of 300 training epochs was reached, or the overall validation loss did not improve for 10 consecutive epochs.

During inference, the overlapping box-wise predictions were assembled to generate the corresponding full-volume outputs. The assembled molecular mask was then converted into a soft mask by iterative morphological dilation. Specifically, the binary mask was dilated outward for 15 successive layers, with newly added voxels assigned cosine-decaying weights from 1 to 0. The resulting soft mask was multiplied with the assembled enhanced density map to suppress residual density outside the predicted molecular region while providing a smooth transition at the molecular boundary. For the predicted local SSIM scores, negative values indicate that the model estimates no reliable positive structural agreement in the corresponding regions. These regions were therefore uniformly treated as low-confidence regions, and the negative predicted scores were clipped to zero.

### Evaluation metrics

The performance of CryoCodex was systematically evaluated on the independent benchmark test set comprising 126 raw cryo-EM density maps, the available half-map data, and 771 structural chains. We used the following evaluation protocol.

#### FSC-0.5

For density map quality assessment, phenix.mtriage^26^ was used to calculate the unmasked map–model Fourier shell correlation (FSC) between each density map and its corresponding reference atomic model. The spatial frequency at which the FSC curve crossed the 0.5 threshold was converted to the corresponding spatial resolution and reported as FSC-0.5. A smaller FSC-0.5 value indicates that map–model correlation is maintained to higher spatial frequencies and therefore reflects improved agreement between the density map and the reference atomic structure.

#### Q-score

the MapQ plugin in UCSF Chimera^28^ was used to calculate Q-scores^27^. Q-score evaluates the resolvability of individual atoms by measuring the real-space correlation between the density profile surrounding each modeled atom and an expected reference density profile. The mean Q-score over all atoms was used to characterize the overall resolvability of the density map, with higher values indicating better-defined atomic density.

#### Model-building performance

To evaluate whether density enhancement facilitates downstream atomic model building, phenix.map_to_model^26^ was used to perform automated de novo model construction on the chain-specific density maps without manual intervention. Prior to model building, local density sub-volumes corresponding to individual chains were extracted from the full density maps using a 4 Å cutoff around the atoms of each chain in the reference model. The automatically generated models were subsequently compared with the corresponding reference structures using phenix.chain_comparison. Protein and nucleic-acid residues were compared using Cα and P atoms, respectively, with a spatial matching cutoff of 3.0 Å. Residue coverage was defined as the fraction of residues in the reference structure that were spatially matched by residues in the automatically built model. Sequence recall was calculated as the fraction of reference residues that were both spatially matched and assigned the correct residue identity. Specifically, sequence recall was calculated as residue coverage multiplied by the sequence-match fraction reported by phenix.chain_comparison.

#### Half-map consistency

To further assess whether the enhanced density features were supported by independently reconstructed experimental data, an independent half-map evaluation was performed for samples with available half-maps. Unmasked FSC curves were calculated for four combinations: between the two unprocessed half-maps, between the two independently enhanced half-maps, and for the two cross-pairings between an enhanced half-map and the unprocessed opposite half-map. Comparison of these inter-half-map FSC curves was used to assess changes in experimental signal consistency after enhancement. In particular, the cross-paired comparisons provide a means of examining whether density features introduced or strengthened in one processed half-map remain supported by the independently reconstructed opposite half-map.

Across comparisons with other enhancement methods, all methods except CryoTEN were evaluated using maps resampled to a voxel size of 1 Å. CryoTEN was applied directly to the native voxel grid of the deposited density maps in accordance with its original processing procedure. Because individual enhancement methods or evaluation tools occasionally failed to produce valid results for specific samples, the mean value for each metric was calculated using only the common subset of samples for which valid results were available for all methods included in that comparison. The common subset was determined independently for each metric. Thus, the number of samples used to calculate the mean could differ slightly among evaluation metrics, while all methods within a given metric were compared on exactly the same set of samples. For comparisons with CryoRes, its output maps were resampled to a voxel size of 1 Å and evaluated within the cubic region determined by the ModelAngelo auto-mask.

## Data availability

The experimental cryo-EM maps and corresponding atomic structures used in this study are publicly available from EMDB and PDB. CryoCodex-generated results for the benchmark dataset are available at https://yanglab.qd.sdu.edu.cn/CryoCodex/.

## Code availability

The source code is publicly available at https://github.com/YangLab-SDU/CryoCodex. To facilitate the use of CryoCodex, a web server has been established on Google Colab.

## Author contributions

J.Y. conceived and supervised the project; B.C. designed and implemented the experiments; B.S. co-supervised the study; All authors reviewed and approved the final version of the paper.

## Acknowledgements

This work is supported by the National Natural Science Foundation of China (NSFC T2225007, T25B2009, 32430063) and the Fundamental Research Funds for the Central Universities.

