## Supplementary Information for "CryoCodex: learning discrete structural representations for cryo-EM map post-processing"

**Supplementary Table 1 | Map quality across cryo-EM density enhancement methods on the 126-map benchmark test set.** FSC-0.5 statistics were calculated on the common subset of 120 maps for which valid results were available for all evaluated methods, whereas Q-score statistics were calculated on all 126 maps. ΔFSC-0.5 and ΔQ-score indicate improvements relative to the deposited maps. Bold values indicate the best performance for each metric.

| Method | FSC-0.5 | Q-score | ΔFSC-0.5 >1 Å (%) | ΔQ-score >0.05 (%) | ΔQ-score >0.08 (%) |
|---|---|---|---|---|---|
| deposited | 6.87 | 0.422 | - | - | - |
| CryoTEN | 4.86 | 0.405 | 27.50 | 0.79 | 0.00 |
| DEMO-EMReF | 4.82 | 0.446 | 45.00 | 33.33 | 8.73 |
| EMReady | 4.83 | 0.450 | 40.00 | 33.33 | 7.14 |
| EMReady2 | 4.62 | 0.460 | 43.33 | 35.71 | 11.11 |
| **CryoCodex** | **3.93** | **0.503** | **75.83** | **73.81** | **57.94** |

**Supplementary Table 2 | Map quality across cryo-EM map enhancement methods on the nucleic-acid-rich subset.** Comparison of the qualities between the deposited and processed maps across a test set of 22 cryo-EM maps with nucleic acid content exceeding 10% in terms of FSC-0.5 and Q-score. FSC-0.5 statistics were calculated on the common subset of 20 maps for which valid results were available for all evaluated methods, whereas Q-score statistics were calculated on all 22 maps. ΔFSC-0.5 and ΔQ-score indicate improvements relative to the deposited maps. Bold values indicate the best performance for each metric.

| Method | FSC-0.5 | Q-score | ΔFSC-0.5 >1 Å (%) | ΔQ-score >0.05 (%) | ΔQ-score >0.08 (%) |
|---|---|---|---|---|---|
| deposited | 5.45 | 0.427 | - | - | - |
| CryoTEN | 4.32 | 0.417 | 25.00 | 0.00 | 0.00 |
| DEMO-EMReF | 4.27 | 0.451 | 35.00 | 27.27 | 4.55 |
| EMReady | 4.42 | 0.448 | 25.00 | 18.18 | 4.55 |
| EMReady2 | 4.02 | 0.474 | 35.00 | 36.36 | 13.64 |
| **CryoCodex** | **3.43** | **0.522** | **80.00** | **86.36** | **68.18** |

**Supplementary Table 3 | Map interpretability of de novo model building.** All atomic models were built using phenix.map_to_model across the entire benchmark dataset. Performance for each evaluation metric is reported separately for the overall dataset (a total of 761 chains) and the nucleic acid subset (comprising 102 chains). Bold values indicate the best performance for each metric.

| Method | Residue coverage (%) | | Sequence recall (%) | |
|---|---|---|---|---|
| | Overall | Nucleic acid | Overall | Nucleic acid |
| deposited | 51.5 | 43.8 | 15.5 | 17.0 |
| CryoTEN | 57.9 | 51.3 | 18.7 | 18.9 |
| DEMO-EMReF | 60.3 | 54.2 | 23.1 | 19.7 |
| EMReady | 64.9 | 57.5 | 25.8 | 23.0 |
| EMReady2 | 66.7 | 64.7 | 25.8 | 23.3 |
| **CryoCodex** | **72.1** | **69.1** | **31.7** | **27.8** |

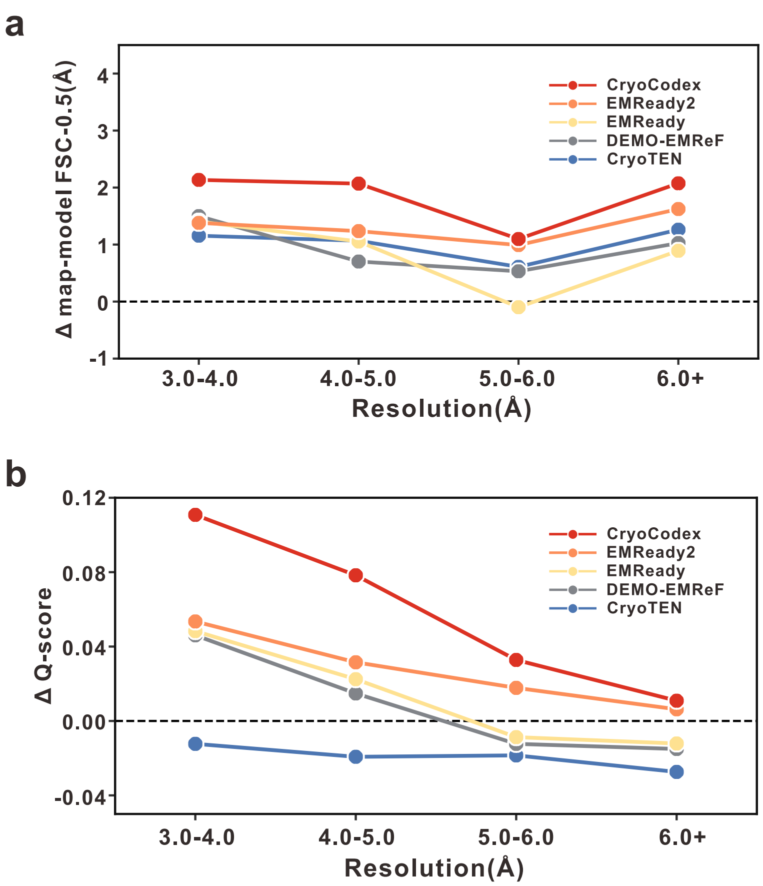


**Supplementary Fig. 1 | Relationship between performance improvements and map resolution for 126 cryo-EM maps. a, b** Average improvements in unmasked map–model FSC-0.5 (**a**) and Q-score (**b**) for various methods across different resolution intervals. The improvements along the vertical axes are calculated relative to the original deposited maps: ΔFSC-0.5 = deposited − method, and ΔQ-score = method − deposited, such that positive values indicate enhanced map quality over the deposited baseline. Higher values represent greater performance improvements for both metrics. For the resolution-dependent FSC-0.5 analysis in (**a**), only maps with valid FSC-0.5 results for all evaluated methods were included. EMD-21225 and EMD-44164 were further excluded because their unusually large deposited map–model FSC-0.5 values would disproportionately affect the apparent trend across resolution intervals, leaving 118 maps for this analysis. The Q-score analysis in (**b**) was performed on all 126 test maps. Full per-map results are provided in Supplementary Data 1.

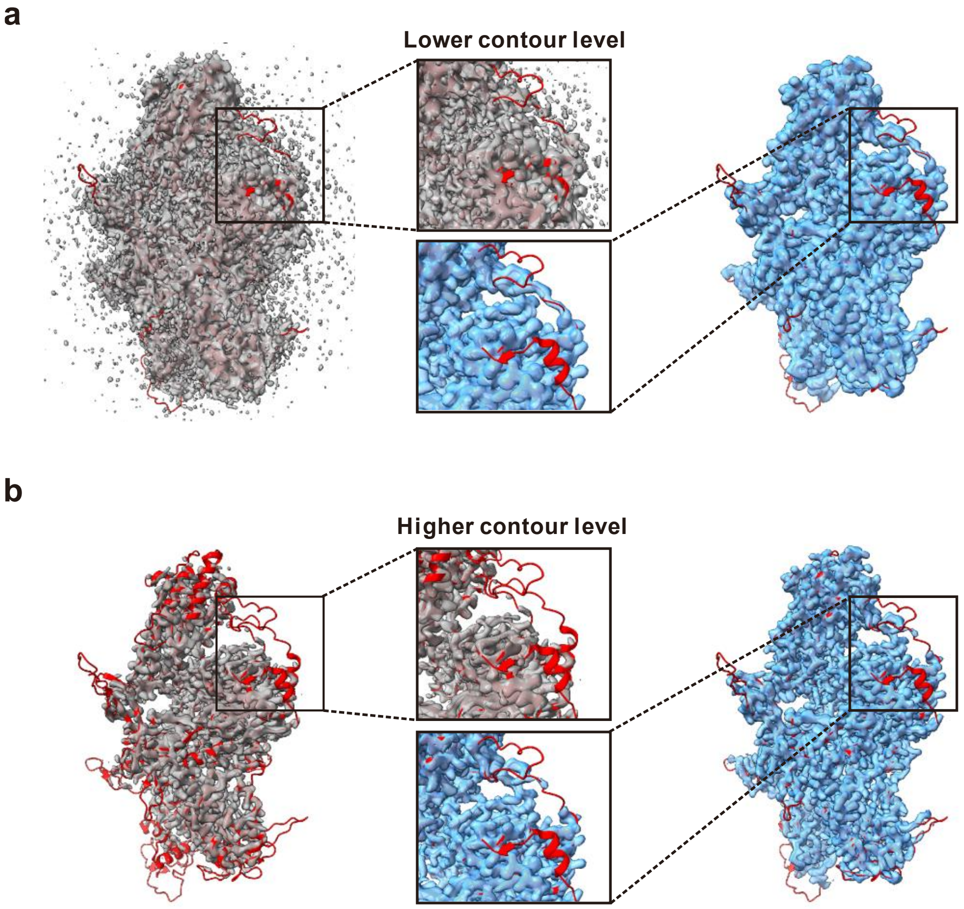


**Supplementary Fig. 2 | Density enhancement in a weak-signal region. a, b** Comparison of deposited (grey) and CryoCodex-processed (blue) maps of EMD-19811 (PDB 8S8P; reported resolution, 3.11 Å) at lower (**a**) and higher (**b**) contour levels. Deposited and processed maps are grey and light blue, respectively; the reference model is shown as a red cartoon. Central insets highlight a peripheral region containing a helical segment and an adjacent flexible loop, where the deposited density is fragmented and close to the noise level. After processing, diffuse background density is reduced, while the weak peripheral density retains its heterogeneous appearance.

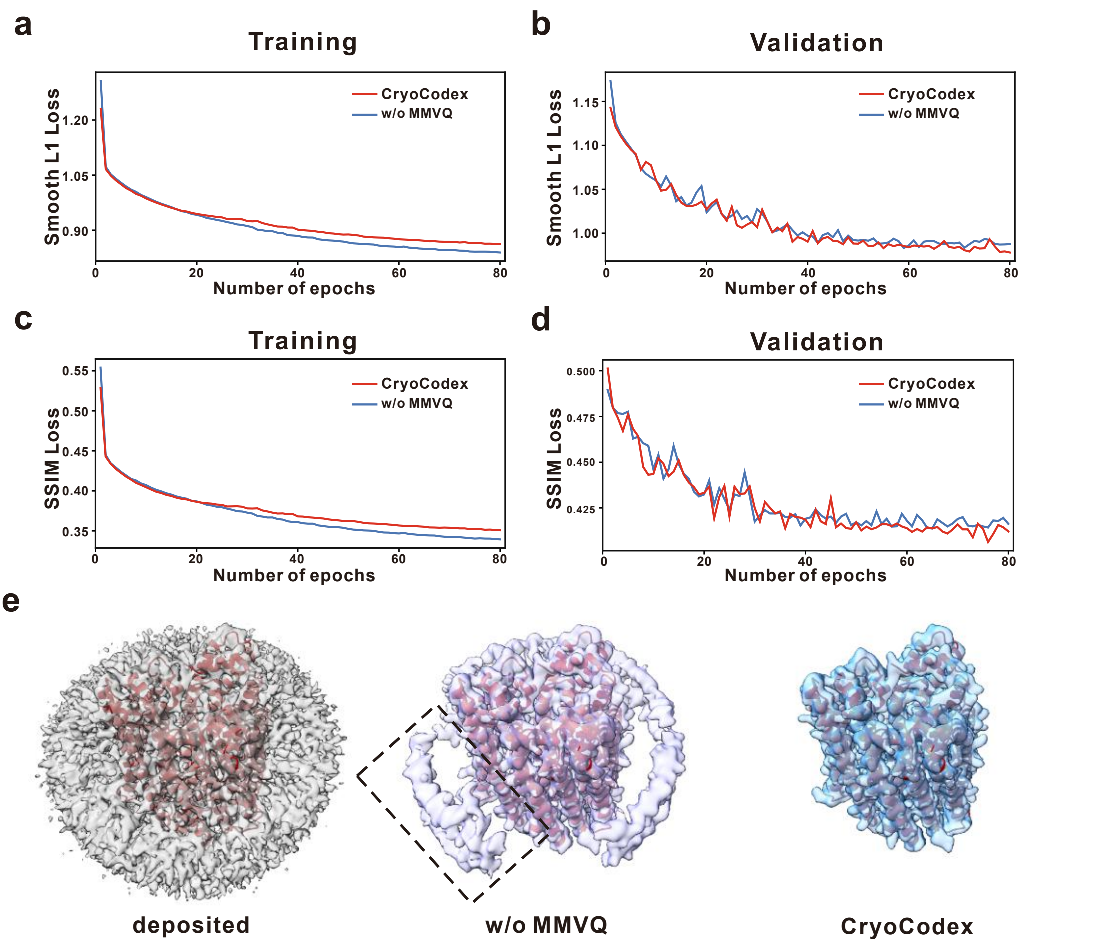


**Supplementary Fig. 3 | Ablation analysis of the MMVQ module in CryoCodex. a, b** Training (**a**) and validation (**b**) curves of the Smooth L1 loss for the density-enhancement task, comparing the full CryoCodex model with the ablated model without MMVQ (w/o MMVQ). **c, d** Corresponding training (**c**) and validation (**d**) curves of the SSIM loss. **e** Representative example of EMD-25135 comparing the deposited map, the output of the model without MMVQ, and the output of the full CryoCodex model. The dashed box highlights residual background density that is more pronounced in the output of the model without MMVQ.

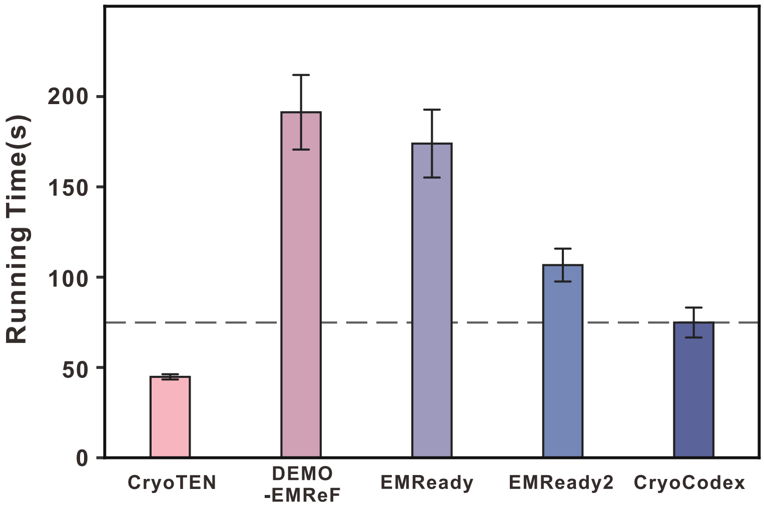


**Supplementary Fig. 4 | Comparison of running time across deep learning-based cryo-EM map post-processing methods.** Running times were measured across all 126 test maps under the same 8 GB GPU-memory constraint. Each method was evaluated using its default inference configuration, with the input batch size set to the maximum value supported under the memory limit. Bars represent mean running times, and error bars indicate the standard error of the mean (SEM). The horizontal dashed line marks the mean running time of CryoCodex.
